# Development of an Ocean Protein Portal for Interactive Discovery and Education

**DOI:** 10.1101/2020.05.29.124388

**Authors:** Mak A. Saito, Jaclyn K. Saunders, Michael Chagnon, David Gaylord, Adam Shepherd, Noelle A. Held, Chris Dupont, Nick Symmonds, Amber York, Matt Charron, Danie Kinkade

## Abstract

Proteins are critical in catalyzing chemical reactions, forming key cellular structures, and in regulating cellular processes. Investigation of marine microbial proteins by metaproteomics methods enables the discovery of numerous aspects of microbial biogeochemistry processes. However, these datasets present big-data challenges as they often involve many samples collected across broad geospatial and temporal scales, resulting in thousands of protein identifications, abundances, and corresponding annotation information. The Ocean Protein Portal (OPP) was created to enable data sharing and discovery among multiple scientific domains and serve both research and education functions. The portal focuses on three use case questions: “Where is my protein of interest?”, “Who makes it?”, and “How much is there?”, and provides profile and section visualizations, real-time taxonomic analysis, and links to metadata, sequence analysis, and other external resources to enabling connections to be made between biogeochemical and proteomics datasets.

## Introduction

For decades, environmental scientists have relied on standard measurements to assess ecosystem change and health, such as temperature, oxygen concentration, nutrient content, chlorophyll abundance and so on.^1-3^ These approaches, while essential in detecting ecosystem level understanding, are limited in their ability to bring about understanding of what each organism within those ecosystems is experiencing and how the organisms respond to environmental change. Recent improvements in “omics” capabilities – consisting of four major omics: genomics, transcriptomics, proteomics, and metabolomics – now allow researchers to begin to open the “black box” of ecosystems to investigate each organism’s catalog of genes (genome), how they choose to deploy those genes in specific environmental settings (transcripts and proteins), and the resulting impact on metabolism and the chemical environment (metabolites).^4-7^ While these new capabilities are exciting, research is still in the relatively early stages of maximizing their utility. Moreover, because every individual biological sample can return thousands to millions of units of raw data (sequence or spectra) these data types are firmly in the realm of big data and bring unique informatic challenges.

We have developed a web portal called the “Ocean Protein Portal” that focuses on developing and improving the delivery of data products related to the measurement of proteins in the oceans, usually referred to as ocean *metaproteomics*. Oceans cover ∼70% of the Earth’s surface and play a critical role in maintaining habitable conditions on the planet. Thus, the continued health of the oceans is an issue of sustainability. Moreover, the ocean and terrestrial microbial communities are responsible for most of the biogeochemical reactions that created and maintain habitable conditions on Earth.^8^ The direct measurement of proteins in the oceans has generated considerable excitement because proteins are the functional units of the cell. They represent where “the rubber meets the road”: enzymatic proteins are the biomolecules that interface with the environment and conduct biogeochemical reactions (Figure 1), rather than the blueprint of genetic potential that genomic data provides. Similarly, while RNA measurements provide information about the transcription of genes, the shorter timescales of RNA production and decay need to be considered in their interpretation. Proteins measurements, with their longer timescales, can be applied as biomarkers of ecosystem health. Additionally, enzymatic proteins that are *directly* responsible for biogeochemical reactions can be measured and their activities estimated to validate global ecosystem models. Individual key proteins have been used to detect specific responses of microbial organisms to nutrients and environmental stressors (e.g. iron, nitrogen, phosphorus, and metabolite transporters)^5, 6, 9-16^ or important biogeochemical reactions (e.g. enzymes that catalyze carbon and nitrogen biogeochemical reactions).^6, 17-20^ As a result, there is a growing interest among experimentalists, observationalists, and modelers to use metaproteomic data for contextual information about their research.

**Figure 1.**
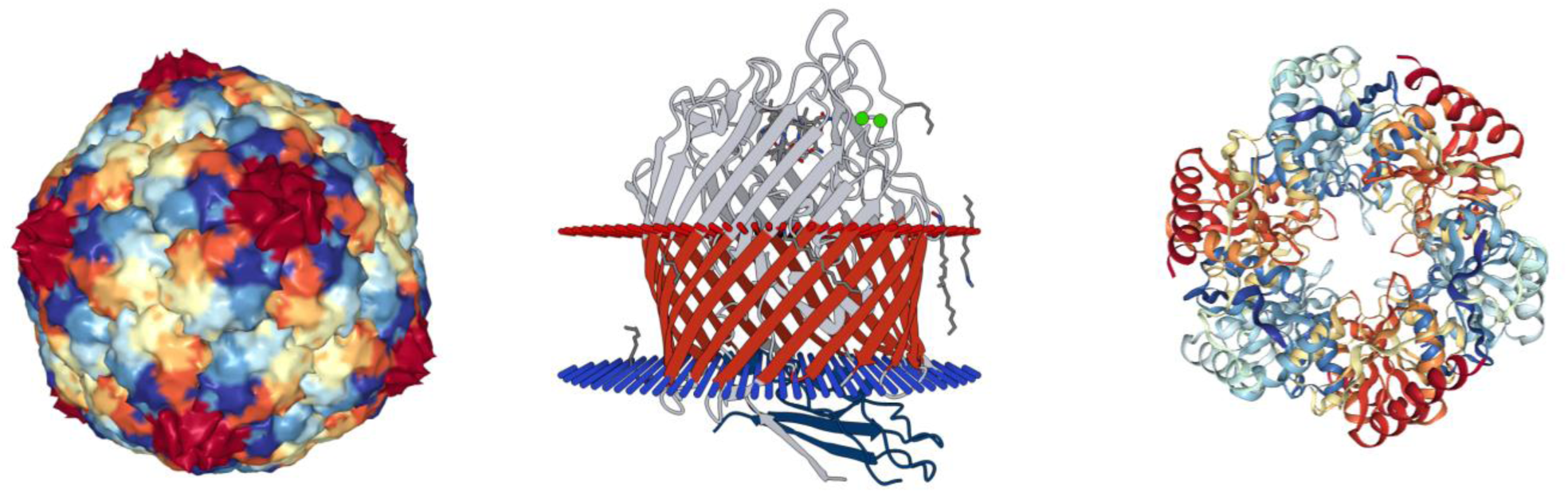
Example 3D structures of common proteins found in the marine environment with important functional roles and routinely found within the OPP. Left: Viral protein capsid of a marine cyanophage.^52^ Center: TonB vitamin transporter spanning the cell membrane.^53^ Right: carboxysome shell protein (CsoS1D) from *Prochlorococcus marinus* MED4 (PBD codes 2XD8, 2GSK, 3FCH).^54^ While genomics shows the potential to make these proteins, protein measurements can show the response of each organism to environmental cues by biosynthesis of specific proteins.

The fields of environmental genomic and transcriptomic informatics is more mature than for protein informatics, with millions of dollars invested to date on data access and analysis portals, including the failed CAMERA project,^21, 22^ the Department of Energy Joint Genome Institute’s Integrated Microbial Genomes and Metagenomes server (DOE JGI-IMG-M),^23^ the Ocean Gene Atlas that uses the Tara Ocean expedition dataset,^24, 25^ and iMicrobe.^26^ In comparison, the Ocean Protein Portal is, to our knowledge, the first investment to date focused on environmental metaproteomic data that produced an operational product in active use across multiple science domains, including oceanography, geobiology, microbiology, and biochemistry communities. Here we describe Version 1 of the Ocean Protein Portal as a means to promote use of ocean metaproteomic data in research across multiple scientific domains and education.

## Results and Discussion

### The Ocean Protein Portal as a Resource to Study Ocean Proteins

The Portal arose from community interest and use case development from the EarthCube ECO-GEO Research Coordination Network focused on environmental ‘omics data. The OPP team represents a collaboration between domain scientists, informaticists, data managers, and computer programmers. The OPP use cases were designed to allow a broad range of scientists and students to discover answers to the questions: 1) “Where is my protein of interest in the oceans?”, 2) “Who makes the protein?”, and 3) “How much is there?”. The OPP is primarily a mechanism to study a single protein query at a time rather than a tool for comprehensive analysis of a metaproteomic dataset. We previously published a metaproteomic viewer that facilitates some metaproteomic data visualization and analysis.^16^ Thus far, the OPP has achieved two milestones in two major categories: the launch of a functioning web user interface (UI) for and essential backend infrastructure for the UI functioning, and educational and outreach activities to promote the study of proteins in environmental settings utilizing the OPP web UI (Table 1).

**Table 1.** Accomplishments to date for the Ocean Protein Portal Project.

|  |  |
| --- | --- |
| Best Practices Workshop and Publication for Data Sharing and Metadata | Saito et al., 2019 |
| Development of Ocean Metaproteomic Viewer python software as test bed for OPP visualizations. Presented and published at Scientific Python conference. | Held et al., 2018 |
| Launch of METATRYP Least Common Ancestor Software and API (used by Portal) | February 2018<br>Saunders et al., 2020 |
| Launch of Ocean Protein Portal Version 1 capable of answering use case questions: 1) Where is protein of interest? 2) who makes it? 3) how much is there? | February 2019 |
| First year metrics over 1000 uses of portal | As of March 2020 |
| Ingestion of protein datasets from Arctic, Antarctic, Pacific, and Atlantic Ocean. Future large datasets expected from Atlantic, Pacific and Antarctic regions, including from BATS and HOT time series stations and the CICLOPS Ross Sea expedition. | Ongoing |
| Use of Ocean Protein Portal in undergraduate and graduate education | Mt Alison College, MIT-WHOI, others |
| Use of OPP for discovery of novel functional protein distributions and publication of data | Mazzotta et al., 2020 |

**Table 2.** File types required by the OPP for full functionality*.

| **File Description** | **File Type** |
| --- | --- |
| Protein Identifications and abundance | **CSV Template** |
| Peptide Identifications and abundance | **CSV Template** |
| Amino acid sequences of identified proteins | **FASTA text file** |
| Metadata expedition and dataset forms | **Rich Text Format (RTF)** |

\* https://github.com/oceanproteinportal/data-file-templates

**Table 3.** Ocean Metaproteomic Datasets Currently within the Ocean Protein Portal.

| Dataset Name | Expedition Number - Location | Filter Type (microns) | Sample | Publication Status |
| --- | --- | --- | --- | --- |
| Metzyme 0.2 | KM1128; Central Pacific Ocean | 0.2-3.0 | 37 | Saito et al., 2014 |
| Metzyme 3.0 | KM1128; Central Pacific Ocean | 3.0-51 | 40 | In Preparation |
| Nunn Arctic-Bering Sea | HLY1301; Arctic Ocean and Bering Sea | 0.003-0.8 | 2 | May et al., 2016<br>Mikan et al., 2019 |
| Morris CoFeMUG | KN192; South Atlantic Ocean | 0.03-3 | 16 | Morris et al., 2010 |
| Walsh Canada Basin | JOIS 2015; Arctic Ocean | 0.2-3.0 | 9 | In preparation |
| Walsh Baffin Bay | ArcticNet2013_CCSG_Amundsen;<br>Arctic Ocean | 0.2-3.0 | 12 | In preparation |
| ProteOMZ | FK160115; Central Pacific Ocean | 0.2-3.0 | 103 | In preparation |
| Ross Sea Net Tow (Bender) | NBP0601; Ross Sea, Southern Ocean/Coastal Antarctica | > 20 | 2 | Bender et al., 2018 |

**Table 4.**
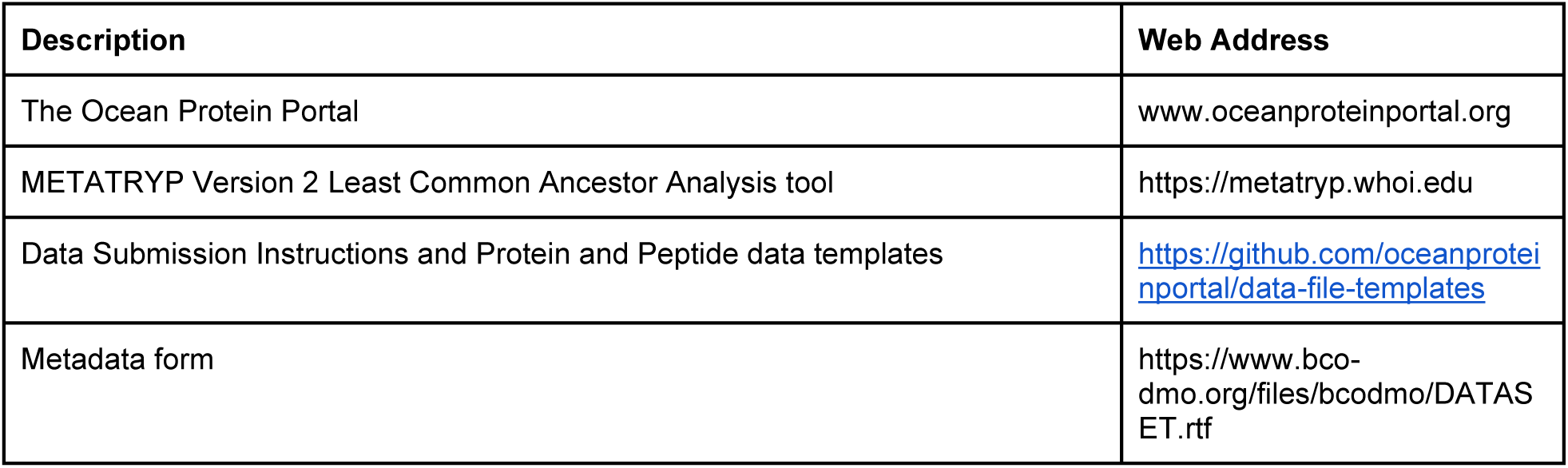
Ocean Protein Portal websites and submission resources.

The OPP UI enables users to answer the three use case questions above for their protein of interest in the oceans via multiple search strategies (Figure 2). The simplest is by entering its common name, for example the carbon fixing enzyme “RUBISCO”, into the “Search Value” text box with the “Search Term” *Protein Name* selected. Wildcard searches (using “*”, for example “carboxy*”) are also allowed since protein names are not standardized and multiple names can be used to describe the same protein in the literature. Alternatively, users can search by using accession numbers of various standardized bioinformatic identifiers, such as KEGG (Kyoto Encyclopedia of Genes and Genomes), UniProt, PFam (Protein Family), or EC (Enzyme Commission number), that allow cross-platform connectivity. Finally, peptide and full protein sequence searches are possible. For full protein sequences, the user enters the protein amino acid sequence, and the OPP breaks the sequence into smaller tryptic peptides – the tryptic peptides being the measured components of the deposited proteomics data – then searches for exact matches of those component peptides in the OPP database. All searches can be narrowed by various parameters (concentration, depth, filter size, dataset, and date range) using the sidebar widgets. Queries return a table of all matches, listing their protein and KEGG names, the dataset and expedition they were identified within, and the quantitative abundance within that dataset (in spectral count units currently). A map of station locations where the queried protein was identified is shown (stations where the protein is found become highlighted; Figure 2), and a map hover over capability provides expedition metadata.

**Figure 2.**
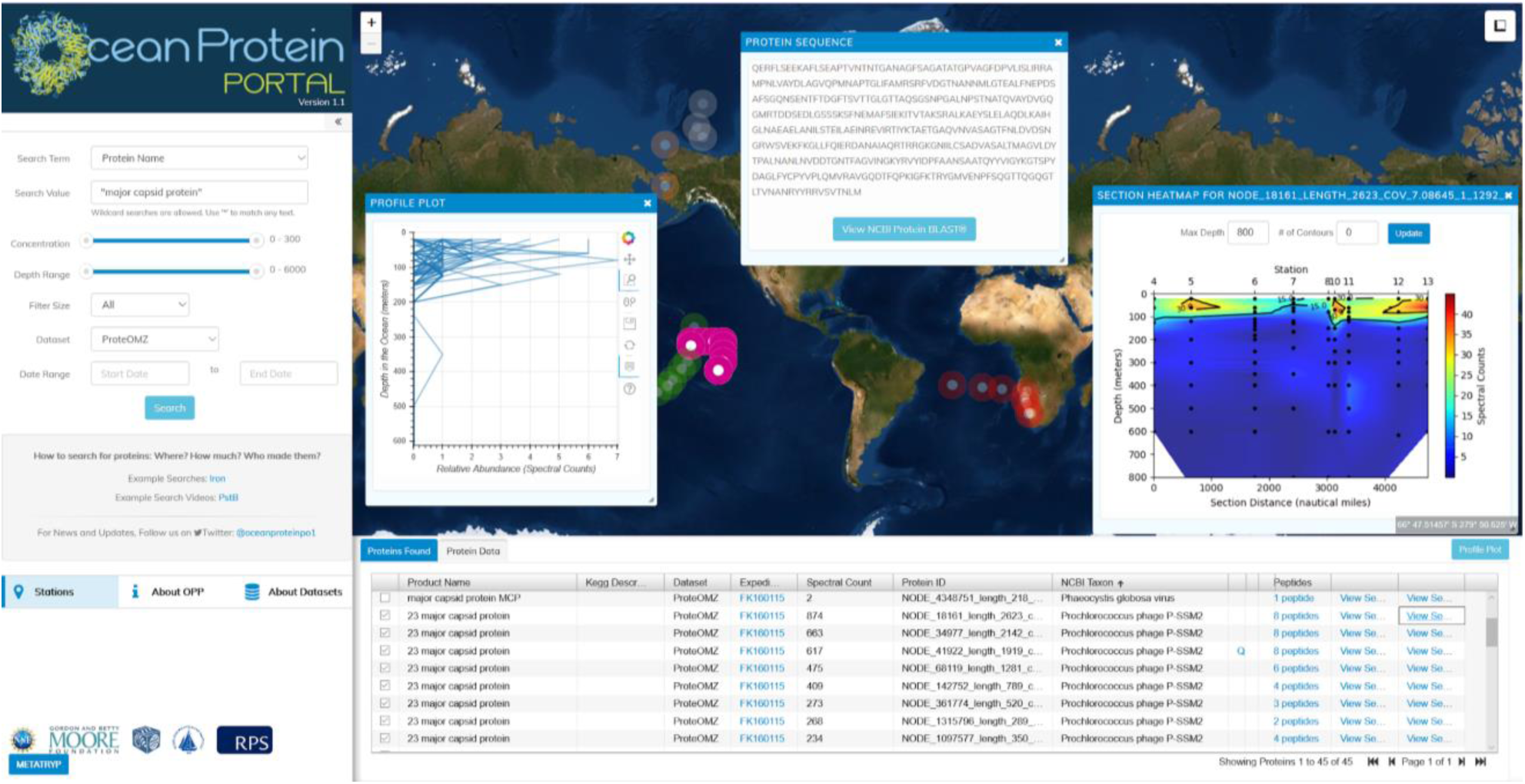
The operational Ocean Protein Portal Prototype. A product name search (“major capsid protein”) showing capsid proteins from marine viruses (Table), vertical profile of capsid proteins (left inset window), protein sequence (center inset) and sectional distribution (right inset) of a major capsid protein from cyanophage, overlaid on the background map of stations (e.g. pink stations). This protein is used to make the physical body of the virus capsid sphere shown in Figure 1 (left), and its distribution across several thousand kilometers of ocean space in the Central Pacific Ocean can be determined with a simple search in the OPP. This protein is one of over 100,000 proteins ingested to date that can be searched for and visualized in the OPP.

After the initial query, users have three options at their disposal for further investigation of their protein of interest. First, users can visualize protein abundance in a vertical profile (1D by depth, “Profile Plot” button) or ocean section (2D by depth with interpolation across transect distance, “View Section” button) mode as pop-up windows (Figure 2). These visualizations use the open source python tools Bokeh and Matplotlib and were prototyped by Held et al., 2018.^16^ Next, users can utilize a suite of links to other bioinformatic resources specific to their protein of interest, leveraging the capabilities of other pre-existing tools. These include BLAST sequence searches (“View Sequence”) that automatically inserts the protein amino acid sequence into NIH National Center for Biotechnology Information’s (NCBI) blastp search box facilitating search of the NIH sequence database as well a hyper-link to the European Bioinformatic Institute’s UniProt sequence database page for the closest related UniProt protein match, when available. The “Expedition” hyper-link routes the user to the full metadata and environmental datasets associated with the sample’s expedition hosted on the ocean environmental data repository at the Biological and Chemical Oceanography Data Management Office (BCO-DMO). Information about datasets and expeditions is also available on the “About Datasets” tab, including contact information of the data generators for each dataset.

Finally, the OPP has a compute capability that enables users to answer “who” is making their protein of interest (Figure 3). This is a key question within the field of metaproteomics because of the possibility that peptide constituents of proteins could be found in multiple organisms present within an individual sample. To address this the OPP utilizes the software tool METATRYP we previously created that searches a database of all tryptic peptides among a group of organisms specified or within meta-omic assemblies from the environment.^5, 27^ METATRYP then identifies peptides that are shared among multiple organisms and reports which organisms share the peptides and calculates the “Least Common Ancestor” (LCA) of the identified taxa possessing this peptide. The METATRYP databases use the NCBI Taxonomy database to identify the ancestral phylogeny of the taxa identified that possess the peptide in question. This analysis happens in real-time using an API call to the “metatryp.whoi.edu” resource. By clicking on the “peptides” link, the user progresses to the “Peptide Found” table, where each peptide component can then be examined for its presence across numerous genomes and metagenome resources. The results can be visualized in heatmap and tree formats to allow the user to gain an immediate understanding of who the “Least Common Ancestor” of the protein constituent is and their associated taxonomic lineages (Figure 3). The OPP is currently using METATRYP Version 2 that has improved performance, can calculate the Least Common Ancestor, and has the capacity to separate its database into genomes, metagenomes, and metagenomic products that is described in a separate manuscript.^27^

**Figure 3.**
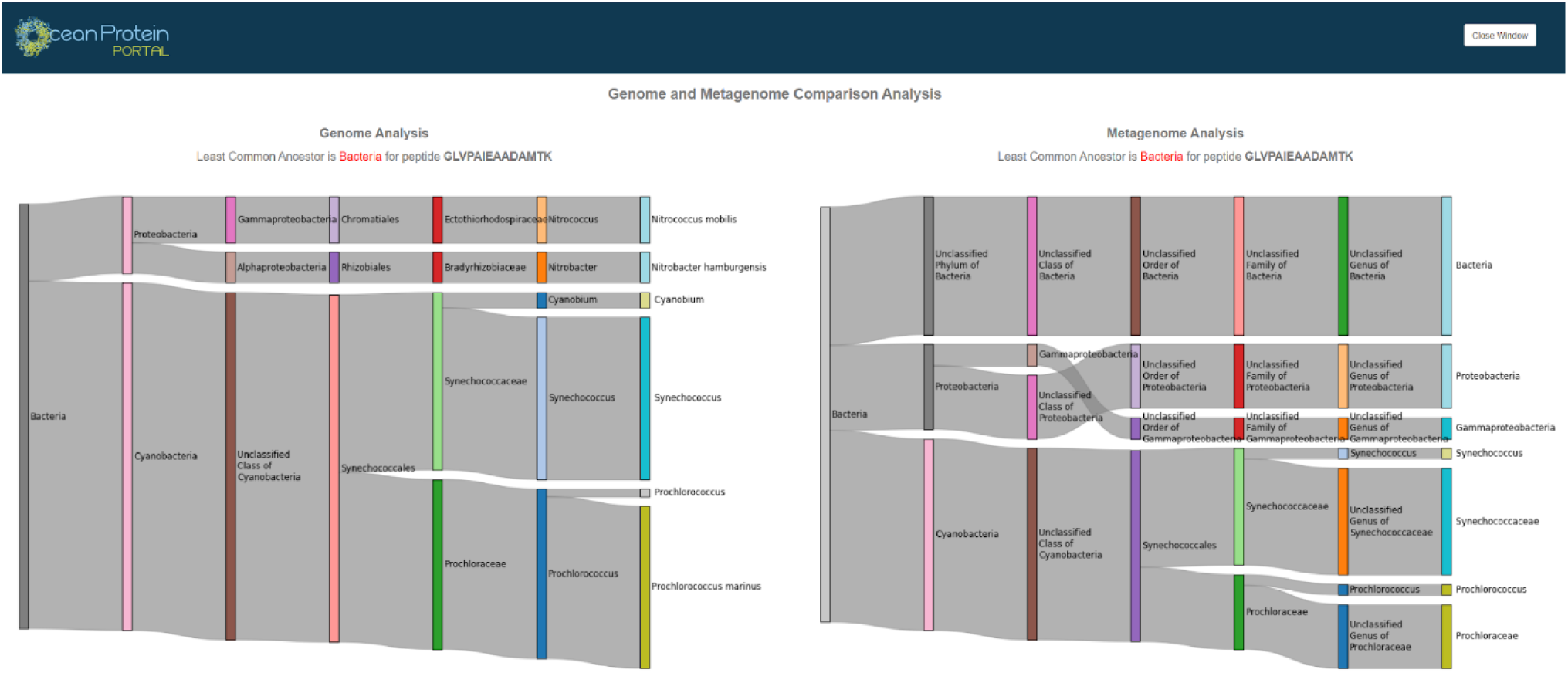
Example of Least Common Ancestor (LCA) analysis representing the taxonomic groups that a queried peptide is found within using the METATRYP API within the OPP UI. This carboxysome shell protein is conserved across multiple Bacterial Phyla, resulting in a similar broad Bacteria LCA level returned using both genomes (left) and metagenome (right) databases within METATRYP.

### Data Ingestion Templates and Data/Metadata Management

Ocean metaproteomic data is not currently standardized in terms of processed output fields and metadata. As a result, the process of ingesting data from a diverse data generator community can be challenging from a data management perspective. Efficient data ingestion is key to sustainability both with regards to recruitment of voluntary data submissions to the OPP by data generators, and in terms of effort needed by data managers and computing staff to successfully ingest data to allow it to function properly within the OPP system.

Through our collaboration with BCO-DMO, we have developed a data ingestion template to facilitate incorporation of complex metaproteomic datasets into the OPP from a variety of data generators with their diverse informatic pipelines. This effort leveraged community driven best-practices that arose from the EarthCube-supported Ocean Metaproteomics Data Sharing workshop.^28^ For every spectral count datapoint, there are an associated 10 metadata fields and 13 annotation fields that can be captured by the current OPP schema. Example metadata reported for each sample includes sampling location (latitude and longitude), depth, date, time, expedition identifier, station number, and filter pore size. Some of these parameters are required, such as the geospatial metadata, while other parameters are optional, such as various annotation fields dependent upon the resolution of the data generators annotation informatics pipeline. We currently do not re-annotate deposited datasets, but hope to add that capability in the future to allow standardized searching across datasets for proteins of interest. To do the database structure is built to allow updated versions with additional supplementary annotation fields that could capture new microbiological and protein function discoveries in previously deposited datasets, while maintaining the data generators’ initial annotations which may link with published research.

### Challenges of Comparisons Across Datasets, Units, and Normalization in the OPP

The current design of the OPP allows users to examine where a protein of interest is in the ocean microbial community, if that protein occurs in at least one of the ingested datasets. One challenge currently is that most ocean metaproteomic data is collected in relative abundance units of spectral counts or precursor intensities (e.g., peptide ms^1^ peak areas), making quantitative comparisons between datasets difficult because of varying instrumentation detection limits and informatic pipelines. The best solution to this is to shift eventually to absolute quantitation of copies of protein per volume of seawater (e.g. fmol/L), that can be compared across space and time with confidence. While absolute quantitation has been used in the ocean, using a technique called targeted metaproteomics,^5, 6, 20^ this datatype is currently scarcer compared to the relative abundance “global” metaproteomics. Moreover, intercomparison and intercalibration of analytical method is needed to validate quantitative values across different data generating laboratories and periods of analysis within laboratories.

Despite these challenges, users will be tempted to compare abundances of their protein of interest across different datasets within the global ocean, comparing different expeditions. While such comparisons may be useful with a binary approach (presence/absence) or relative quantitation approach, we have cautioned users from meta-analyses. Instead, we and encourage users to contact data generators, and if appropriate to collaborate with them on interpretation of results to avoid misinterpreting data as explained in the OPP data use policy found below and on the “About OPP” page in the UI.

Normalization of data is also a factor to consider in the interpretation and comparison of results. While the OPP does not currently have any stipulation as to the type of spectral count unites being used, we encourage the use of non-exclusive total (un-normalized) spectral counts to avoid poor search query performance and/or limit distortion of marine vertical community structure. This rationale, which is specific to metaproteomics and ocean vertical profile sampling, respectively, is explained here in more detail. For background, a spectral count is an easy to calculate unit defined as the count of mass spectra with a peptide identified within it. Within each sample analysis, 10’s of thousands of spectra are typically collected, and spectra that match to a peptide from proteins predicted by a specified sequence database are tabulated by peptide-to-spectrum mapping (PSM) algorithms. Software that calculates spectral counts often have the ability to calculate normalized spectral counts; for example, one normalization strategy is where each protein within the dataset is divided by the total number of spectral counts within the sample and multiplied by the average spectral counts in all samples. These normalizations can be problematic in metaproteomics samples because the number of PSMs and resultant total spectral counts can vary greatly between sampling sites and times as large changes in biological community structure occur. This decrease in total spectral counts may be due to limitations of the database being used with fewer peptide identifications with depth, or increased interferences by organic molecules and degraded peptides that are known to be prevalent with depth.^17, 18^ An example of this problem is shown in Figure 4 where data dependent analysis global proteome analyses of microbial biomass samples from 20m to 800m depth in the North Atlantic Ocean, all injected with a uniform amount of protein onto the LC-MS, nevertheless results in more PSMs observed in the shallower waters (shown by greater sum of total spectral counts) where microbial biomass is more abundant and better characterized by metagenomic databases. Four representative microbial proteins that have maxima at different depths show how normalization can cause considerable biases in their vertical structure. Surface proteins (UrtA1 and TufA) tend to be less abundant and deep proteins are more abundant (GroEL and OpuAC) than the comparable total spectral counts at each depth due to normalization. Based on these biases, it is not obvious that this type of normalization provides any benefit to the analysis. Alternatively, normalization to total protein extracted with depth may be more useful to realistically portray protein distributions (Saunders et al., in prep).

**Figure 4.**
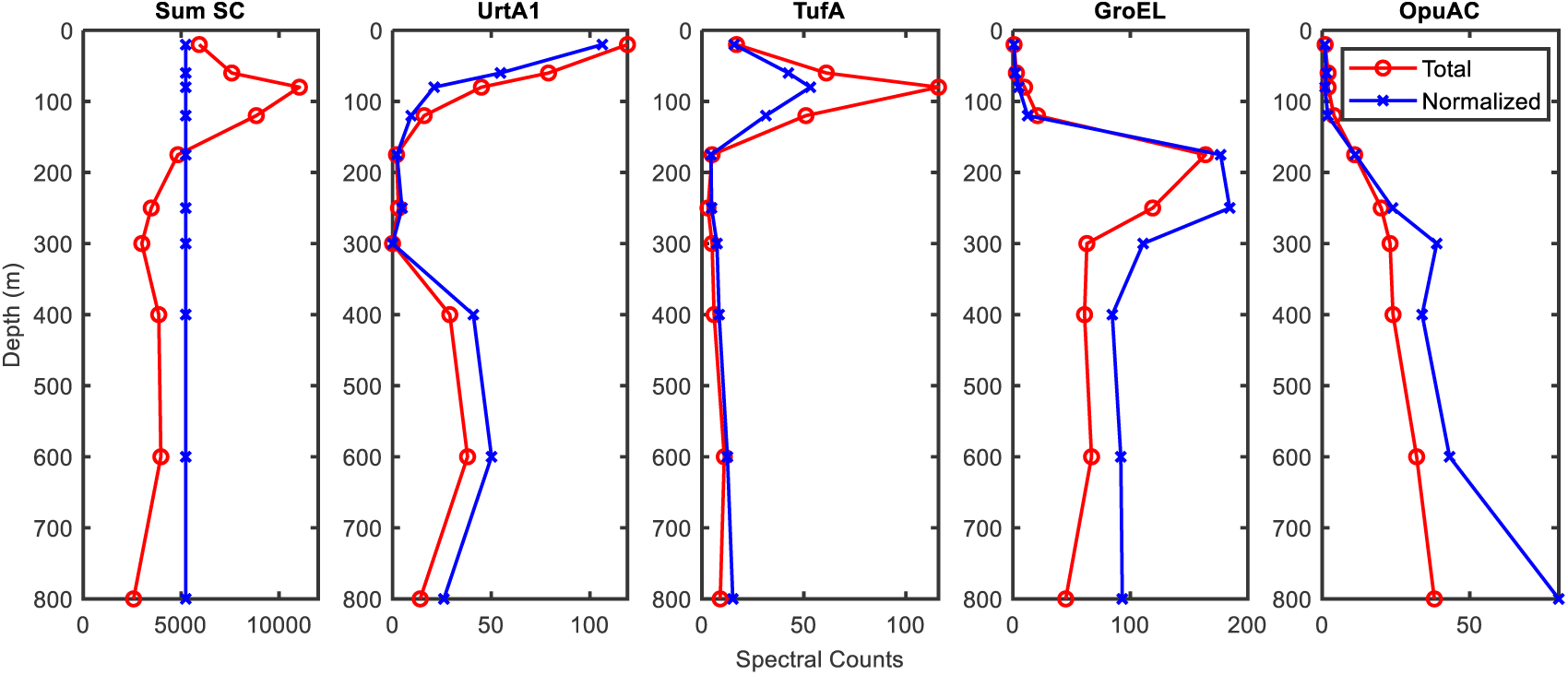
Normalization biases in metaproteomic data across depth in the ocean at the Bermuda Atlantic Time Series Station (31°40′N 64°10′W) in the North Atlantic Ocean on April 14, 2018 collected on 0.2 μm filters by Clio AUV. Left: Sum of total spectral counts (SC) for all proteins at each depth (red circles) and sum of spectral counts after normalizing to the average of all samples (blue crosses). Profiles for four microbial proteins that are abundant at the surface (urea transporter UrtA1), chlorophyll maximum (elongation factor TufA 80m), mid-depth (chaperonin GroEL at 175m), and deep (ligand binding protein OpuAC family at 800m). Changes in the biological community result in greater numbers of peptide-to-spectrum matches in the upper water column. This creates biases when normalization is conducted across depths by treating them as “similar” biosamples, with decreased shallow and increased deep normalized counts compared to the total counts. Data from Breier et al., submitted.

The NSAF normalization (Normalized Abundance Spectral Factor) and similar approaches (APEX; emPAI) that take into account protein length are also often used to prevent bias towards the identification of large proteins with many tryptic peptides over shorter protein sequences with fewer tryptic peptides.^29-31^ These corrections seem logical in laboratory experiments, but the metagenomic and metatranscriptomic databases that spectra are mapped to are often replete with incomplete open reading frames, resulting in incorrect molecular weight estimations and the resulting length corrections to be incorrect. Hence we currently discourage NSAF units within the OPP, at least until the use of newer metagenomic assembly techniques becomes more widespread, such as when PSM solely to metagenome assembled genomes (MAGs) and single amplified genomes (SAGs) is routine.

Finally, there can be calculations of exclusive spectral counts, where each spectrum is only allowed to map to one sequence within the database, even if that peptide sequence is found within multiple proteins from the PSM search database. The occurrence of a peptide within multiple metagenomic or metatranscriptomic reads is a common occurrence within metaproteomics as the natural diversity found within the environment can be captured with sequencing, resulting in multiple sequence assemblies that have both high sequence identities and share identical tryptic peptides. Software such as Scaffold by Proteome Software allows output of “exclusive” spectral counts where spectra of peptides are restricted to map to only one protein sequence through use of a straightforward parsimony algorithm where the protein that has the most peptide matches captures those spectral counts, or alternative “total” spectral counts where those peptide spectra are allowed to map to multiple proteins simultaneously. In cases where a meta-analysis of an entire dataset is being conducted for overall protein taxonomic diversity or function, use of exclusive spectral counts are important to avoid double counting peptides. In contrast, in the single protein-query use case that the OPP is built for, allowing sharing of those peptides can actually be important in allowing exploration of the diversity of protein sequences that exist because exclusive spectral counts can “rob” the peptides from alternate near identical protein sequences that may also be present, potentially suppressing the identification of rarer proteins in these communities. While a future update of the OPP could facilitate switching between multiple unit types (e.g. total, exclusive, and normalized to total protein spectral counts), it is nonetheless important to articulate the implications and pitfalls of each approach in dealing with complex metaproteomic dataset. While emerging targeted metaproteomic data in absolute abundance units can avoid many of these normalization and attribution problems, the ease with which relative abundance datasets containing thousands of proteins (in spectral counts or peak intensities) are generated makes them attractive to broad audiences for hypothesis generation and discovery, and hence the OPP is designed to serve this datatype.

### The OPP Schema

An initial data description (schema) for the OPP was generated along with the OPP prototype using a Resource Description Framework (RDF) format as an extension from the BCO-DMO schema.^32^ This Ocean Protein Portal Data Type Schema (OPP-DT)^33^ defines the different observational entities (e.g. peptide spectral counts, protein spectral counts, FASTA sequence), the associated metadata entities (cruise, sampling date, depth of sample, etc.), and the basic relationships between these entities currently in the portal. Figure 5 illustrates the OPP-DT subclass *Total-Spectral-Counts*, the observational entities within this class, the associated metadata entities, the relationship requirements between these entities, and an example of where a specified metadata entity can be linked out to other scientific data catalogs. This database schema allows for functioning of the OPP web application UI. Additionally, this schema facilitates submission of data into the OPP and help users of the OPP interact with the data through a clear understanding of the relationships between the data fields.

**Figure 5.**
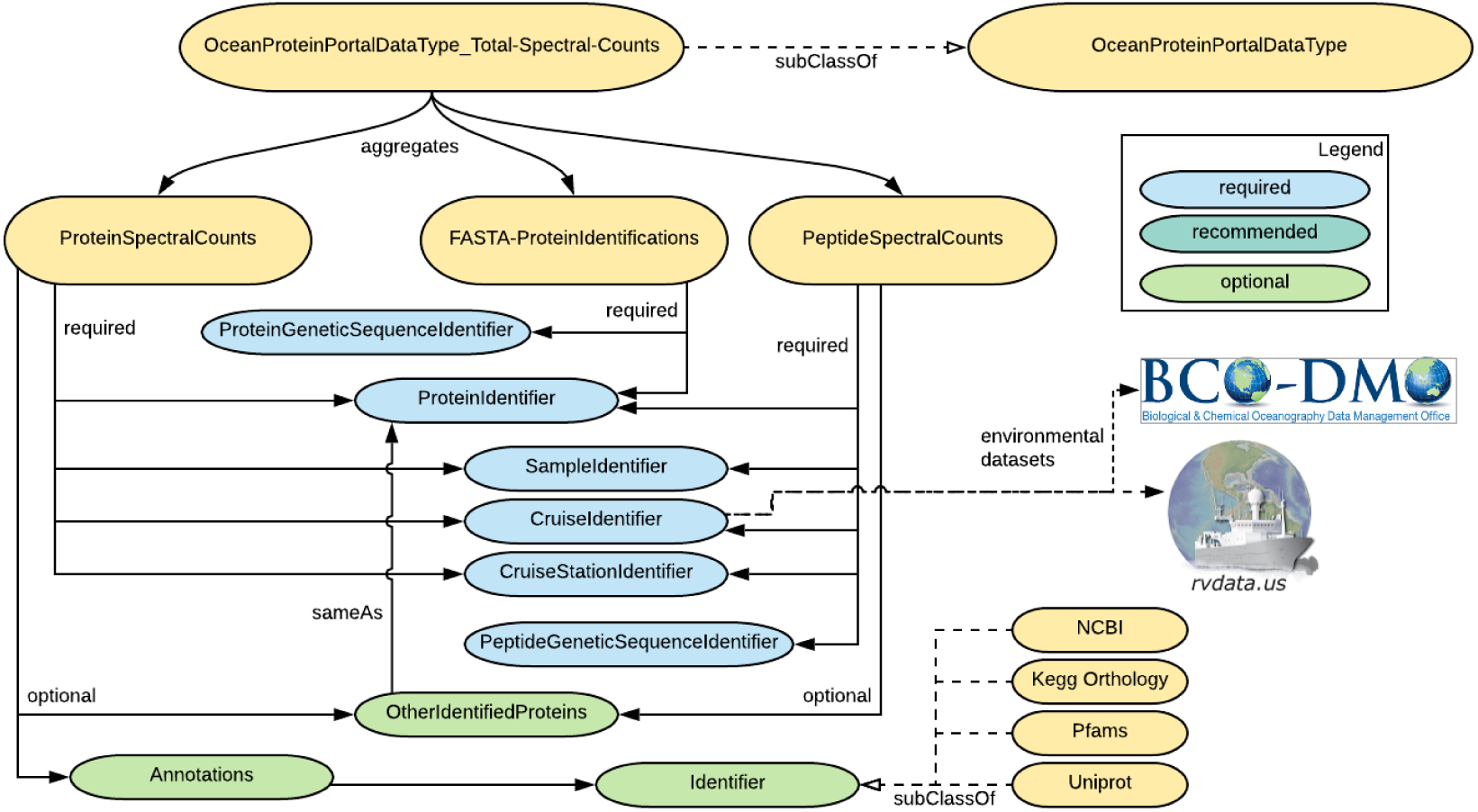
Identifier relationships from the Ocean Protein Portal Total Spectral Count Data Type (OPP-DT) ^33^. Illustrates the various relationship requirements between the three aggregate data types that comprise an OPP Total Spectral Count dataset.

### Technical Aspects

The OPP Version 1.1 is currently built using an Elasticsearch database for protein and peptide data, that is accessed by the UI, generated with Django, Javascript, OceansMap, Bokeh, and Matplotlib code. METATRYP Version 2 Least Common Ancestor software uses PostgreSQL and Python.^27^ Ingestion of data occurs through a process where data generators deposit data for the three file types described in Table 1 according to specified data templates while working with a BCO-DMO data manager. Complete research expedition metadata and co-located environmental datasets are discoverable through the BCO-DMO project pages (linked from the OPP). Both OPP and METATRYP are hosted on servers at the Woods Hole Oceanographic Institution.

An ingestion pipeline has been developed through the application of metaproteomic domain-specific data templates into Elasticsearch using custom scripts and Minio file storage, and has been tested within the BCO-DMO informatics ingestion and data management pipeline. This ingestion pipeline approach utilizing specified templates eases the database relationship connections in Elasticsearch among the data fields names in accordance with the specified OPP ontology. We also used the Frictionlessdata data package to link the three files together which can be expanded upon for further development of the OPP. The ontology design for processing these datatypes follows the Research Data Alliance output and recommendations from their Data Type Registry Working Group.

### Data Use Policy

The OPP is adopting the data use policies similar to the GEOTRACES program, where correct attribution and citation is viewed as an important aspect of the data policy. Moreover, the 2017 Workshop participants for Best Practices in Data Sharing^28^ recommended that users interested in using metaproteomic data sets in publications contact data generators and consider discussing collaboration if using their metaproteomic data. This serves two important purposes: First, there is a danger that non-expert users may misinterpret or misuse data resulting in incorrect interpretations given the youth of the metaproteomic data type especially when considering issues of cross dataset comparisons and normalizations. Publication of interpretations made from incorrect data use could damage broader community confidence in the metaproteomic data type. Second, attribution to, and collaboration with, the data generators will create a valuable incentive for data generators to share future datasets in the OPP’s data search and visualization environment, versus solely depositing data in raw spectra repositories where the data will not be accessible to broader communities outside of proteomics. Hence, the data policy outlined here is useful to the sustainability of the OPP. We anticipate that use of visualizations in publications generated from the OPP could become commonplace and upon publication of the original datasets could occur with simple citation and/or permission of the data generators.

### OPP Scoping Decisions

The OPP was scoped to allow it to be launched within a short time window, to avoid becoming obsolete by tying itself directly to specific proteomic informatic pipelines, and to be lightweight computationally and in terms of code maintenance in order to control upkeep costs for long-term sustainability. A key decision made thus far was for the OPP to accept processed protein and peptide data from depositors, rather than raw mass spectral data. The OPP does not conduct computationally expensive spectral-level re-analyses. These scoping decisions are also important in allowing the domain expert data generators to select and develop their preferred informatic pipelines. There are many up-stream proteomic pipelines used by data generators that produce comparable results, including the peptide-to-spectrum mapping search engines Sequest, Comet, X!Tandem, Mascot, MS-Fragger and OMSSA etc.; Data Independent Acquisition (DIA) and targeted search tools including Skyline, DIA-Umpire, Scaffold-DIA, and EncyclopeDIA etc; and multiple validating and integrating proteomic data systems such as Scaffold and the Trans-Proteome Pipeline.^34-46^ The OPP aims to leverage these packages by accepting the processed data produced by whichever package the data generator utilizes. The OPP was designed to accommodate versioning of submissions and associated metadata to enable data producers to make improvements to their pipelines and update datasets through the OPP data management in collaboration with BCO-DMO. Raw spectra repositories are available through the ProteomXchange, datasets deposited to the OPP can be linked to these repositories allowing expert users to conduct their own re-analyses if they choose to. Finally, the OPP is also not a metagenomic or metatranscriptomic portal given the large amount of resources previously dedicated to those datatypes described above, but can connect with them through hyper-links currently, and perhaps directly in the future using APIs.

### Metrics to Date

The OPP is an online tool launched in 2019 and is in active use. Since its launch, the OPP has ingested and is serving 8 large metaproteomic datasets from multiple data generator laboratories and each dataset can have multiple stations and depths within it. Data are from the Atlantic,^10^ Pacific,^6^ Arctic^17, 18, 47, 48^ and Antarctic (Ross Sea)^14^ Oceans totaling 220 samples, containing 108,549 proteins and 1,581602 peptides altogether. Note this is roughly equivalent to the number of samples within the well-known Tara Metagenome project.^49^ In parallel, the Least Common Ancestry software METATRYP (Version 2) is operational as a standalone tool and also is connected to the OPP via an API, and contains a total of 182,354,079 unique peptides within the database from 142 genomes, 3 metagenomes, and 4,782 specialized genome assembly products (MAGs and SAGs) to date. Use metrics from Google Analytics include over 1300 website use instances of the OPP to date by 700 unique users (Figure 6, left), publication of protein distribution patterns and visualizations from the OPP.^50^

**Figure 6.**
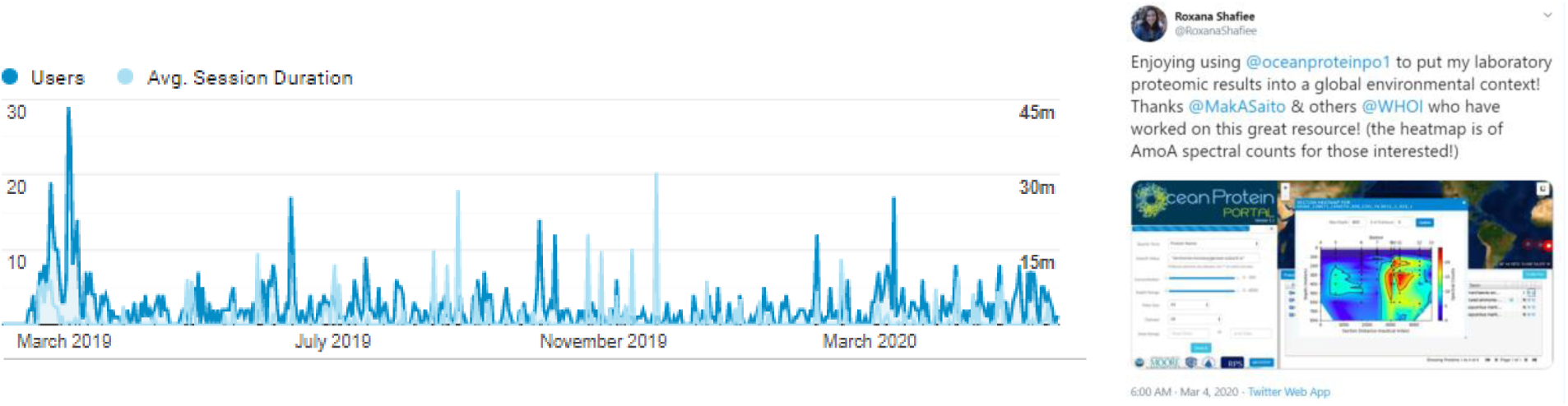
Left: Users and average session duration metrics for the Ocean Protein Portal to date, with unique users totally ∼700 since the launch in Spring of 2019. Right: Social media feedback from a graduate student at Oxford University UK.

### Sustainability

As with all data portals, the OPP faces challenges in operational sustainability and the development of improvements to increase functionality. It was designed with sustainability in mind, by minimizing expensive real-time compute capabilities, by leveraging open-access software, limiting the scope of data types accepted into the OPP, and not attempting to conduct real-time spectral analysis. The current funding model is to use grants for feature development, and “Broader Outreach” funding within core oceanography grants for operational costs (virtual machine, storage, data ingestion, code maintenance). Critical to this effort is for ingestion efforts to be streamlined through the data templates and ingestion pipeline described above to be sufficiently lightweight in data management conducted in collaboration with BCO-DMO.

### Educational Use

In addition to the use in research, we hope the OPP will be a useful tool in education. The OPP can provide students a means to understand how the otherwise invisible molecules they learn about in biology and chemical classes are deployed by life in the natural environment. For example, students can observe how enzymes involved in carbon fixation and photosynthesis are concentrated in the upper layers of the ocean where light penetrates. There has already been interest in educational use of the OPP. For example, the portal is being used in undergraduate teaching and thesis research projects at Mount Allison University (Amanda Cockshutt, pers. comm.) and within graduate microbiology, marine bioinorganic chemistry, and marine microbial biogeochemistry courses. Finally, there is an active social media account that has helped to generate interest and traffic to the OPP, as well as facilitate communication between users and the development team (Figure 6, right). Future curriculum development could help enable teachers and professors in using the OPP.

### Future Improvements

A number of future improvements are planned if resources can be acquired. The current sequence-based search capability of the OPP allows the user to interrogate the dataset independently of annotation information, and hence is useful in situations where the protein function is not yet known or well-characterized, as is the case for many nutrient transporters. Currently, sequence search sends full length sequences to the METATRYP API which digests the sequence into predicted tryptic peptides, then searches them against the OPP peptide database. While this search avenue is operational, it often does not produce any search results because the OPP requires identical string matches of the query peptide against peptides in the OPP database for identification, and hence does not provide flexibility for sequence variability associated with natural biological diversity that users are accustomed to from standard sequence alignment tools (e.g. BLAST: Basic Local Alignment Search Tool^51^). In the future, we hope to incorporate a BLAST-like search of query sequences against peptides in the database allowing for some sequence variability to exist between the user’s query sequence and the OPP database peptides.

## Conclusions

The Ocean Protein Portal was developed to facilitate research and education by allowing users to search for a protein of interest, and examine its distribution in nature. Moreover, taxonomic assessment of the protein is enabled through the use of Least Common Ancestor analysis. With growing interest in ocean health, the OPP could be a valuable resource in connecting a broad audience to ocean metaproteomic datasets, enabling greater understanding of ocean biochemistry and how global and regional environmental change is influencing these critical environments.

## Notes

The example spectral count dataset in Figure 4 was described and submitted in a prior manuscript (Breier et al., submitted), and its corresponding mass spectrometry files have been deposited to the PRIDE Archive under project number PXD018067.

## Acknowledgements

The development of the OPP was supported by an NSF EarthCube grant: “Laying the Foundation for an Ocean Protein Portal” (NSF 1639714), and as part of the broader impacts of NSF-OCE grants 1850719 and 1658030. The underlying METATRYP peptide taxonomic software was developed in a grant from the Gordon and Betty Moore Foundation Marine Microbiology Initiative program (GBMF #8453). JKS was supported by a NASA Postdoctoral Fellowship. The OPP team is a collaboration between the Saito laboratory, the Information Services Application group, and the Biological and Chemical Oceanography Data Management Office all at the Woods Hole Oceanographic Institution. Consulting services were provided by the RPS group. The efforts of the participants of the Data Sharing Workshop for Ocean Metaproteomics (May 2017) were also instrumental in developing best practices for ocean metaproteomics data sharing. Finally, we are indebted to the data contributors described in Table 3 without whom the Ocean Protein Portal would not exist.

